# Human Milk Oligosaccharides Modulate Inflammatory Responses and Lipid Metabolism in a Human Intestinal Organoid Model

**DOI:** 10.1101/2025.03.04.640785

**Authors:** Ellie Slater, Jaesub Park, Thomas Dennison, Tim Mak, Igor Bendik, Ateequr Rehman, Bernd Mussler, Frank Wiens, Komal Nayak, Matthias Zilbauer

## Abstract

Human milk oligosaccharides (HMOs) are a major component of human breast milk and have significant protective effects on infant gut health. HMOs are also present in the amniotic fluid surrounding the developing fetus in the womb. Beginning at 10 weeks of gestational age, HMOs enter the fetal gut through the swallowing of amniotic fluid. The effects of this earliest exposure to HMOs on the gut epithelium, which pre-term infants partially miss, remains unknown. In this study, we investigate the effect of a blend of HMOs, including 2’-fucosyllactose (2’FL), 3’-sialyllactose (3’SL), and difucosyllactose (DFL), on intestinal epithelial cells, examining their role in steady-state conditions and during inflammation. By utilizing advanced intestinal epithelial organoid models derived from fetal and pediatric donors, we reveal developmental-stage-specific responses to this HMO blend, as well as its potential to mitigate inflammatory damage. These findings offer important insights into the role of HMOs in early-life nutrition and gut health, lending compelling evidence to support their inclusion in infant nutrition strategies.

## INTRODUCTION

Human milk oligosaccharides (HMOs) are a diverse family of over 200 molecules, comprising the third most abundant component of human breast milk after lactose and lipids [1,2]. The well-documented health benefits of breast milk and advancements in HMO synthesis have fuelled growing interest in their potential applications as nutritional supplements and therapeutic agents.

Current research predominantly focuses on the role of HMOs on gut health during the neonatal period, where they are introduced orally through breastfeeding. Several studies have demonstrated that HMOs affect gut health indirectly by modulating the gut microbiome, shaping its composition and colonization. Furthermore, emerging evidence suggests that HMOs may also affect immune cell trafficking in the intestine, indicating a possible role in immune homeostasis and inflammation [3–9].

While these studies highlight the indirect effects of HMOs on neonatal gut health, the direct impact of HMOs on intestinal epithelial cells (IECs) remains largely unexplored. Limited research, mostly from murine models and cancer cell lines, suggests that HMOs can promote epithelial cell maturation and enhance barrier function [10–12, 13–14]. However, these models often fail to reflect human physiology or the specific developmental context of the intestinal epithelium. Furthermore, although HMOs have been proposed to influence immune responses in the gut, their direct impact on IECs during inflammation is unknown.

Before breastfeeding, early exposure to HMOs occurs during gestation through the amniotic fluid. At 10 weeks of gestation, the fetus begins swallowing the amniotic fluid, resulting in enteral exposure to various bioactive factors, including HMOs [15–18]. By full term, the fetus swallows approximately 500 mL of amniotic fluid daily [19]. This early exposure represents a critical, yet poorly defined, window in which HMOs may influence fetal intestinal development. The fetal intestine is distinct from the neonatal gut, characterized by the absence of gut microbiota [20] and an intestinal transcriptome indicative of high stem cell activity and differentiation [21–22]. By week 10, intestinal villi and crypts begin to form and fully differentiated secretory goblet cells emerge by week 12 to establish a protective mucus layer critical for barrier function and adaptive immune regulation [23]. During this developmental period, we hypothesize that HMOs may direct the fetal intestinal epithelium to an optimum trajectory for organ development and provide resilience against common challenges, including infections, inflammation and environmental insults [24–29].

Research on fetal gut biology has been limited by various ethical and technical constraints, as well as a lack of physiologically relevant models. The development of human mucosa-derived intestinal epithelial organoids (IEOs) has provided unprecedented opportunities to study human gut biology. IEOs, derived from adult tissue-resident stem cells, faithfully recapitulate intestinal epithelial structure and function, including tissue and developmental stage-specific features [30–32]. Importantly, IEOs allow for controlled studies of epithelial cell-specific responses under both steady-state and inflammatory conditions, independent of confounding factors such as immune cells or the microbiota.

In this study, we utilize IEOs derived from fetal and pediatric donors to examine the effect of an HMO blend on epithelial cell function. We evaluate the effects of these HMOs on IECs under steady-state and inflammatory conditions, revealing differential effects of HMOs in the presence of inflammation. Additionally, we address distinct developmental stage-dependent responses to the HMO blend, hypothesizing that HMOs could play a critical role in shaping intestinal development and resilience against early-life challenges. Our findings provide novel insights into the role of HMOs in early-life nutrition and gut health, emphasizing their importance even before birth.

## RESULTS

### Exposure to the HMO Blend Results in Improved Barrier Function of Human Fetal and Pediatric IEOs

Human intestinal epithelial organoids retain key features of their tissue of origin, accurately reflecting the developmental stage of the patient. Using a previously established method, we derived organoids from fetal and pediatric tissues to investigate the effects of various HMOs on the intestinal epithelium at two critical developmental stages (Fig. 1a).

**Figure 1.**
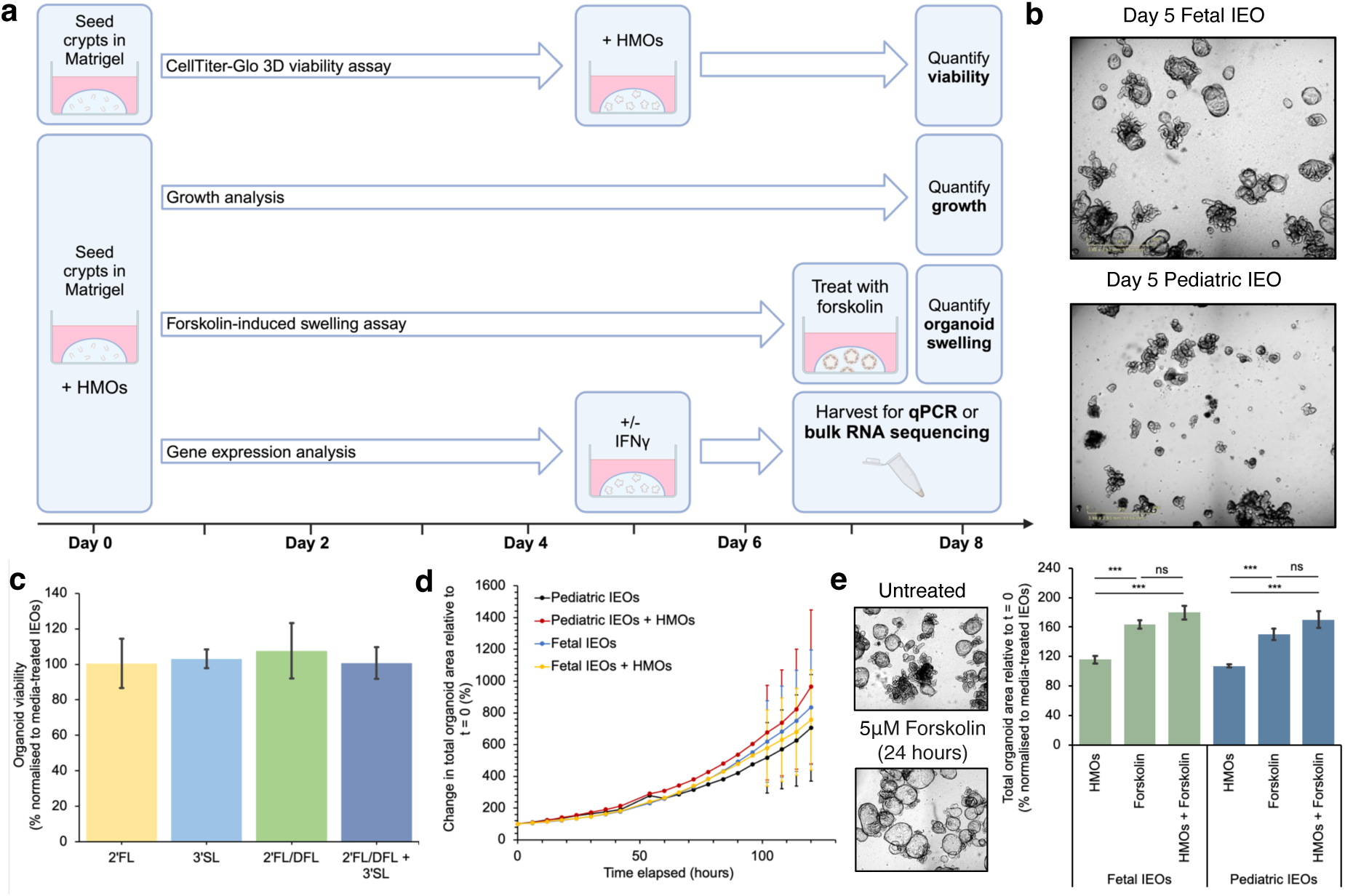
Human milk oligosaccharides (HMOs) support fetal and pediatric IEO growth, viability and barrier integrity under steady-state conditions. **(A)** Illustration of the strategy to test the effect of HMOs on human small intestinal organoids. **(B)** Standard morphology of fetal (top) and pediatric (bottom) intestinal epithelial organoids (IEOs) after five days of growth under the described conditions. **(C)** Organoid viability after basolateral stimulation with HMOs from days 5**-**7, measured using the CellTiter-Glo 3D viability assay and expressed as percent viability compared to media-treated control wells (n = 3). Pediatric organoids were treated with four HMO combinations: 1) 2-fucosyllactose (2’FL), 2) 3-sialyllactose (3’SL), 3) difucosyllactose (DFL) and 2’FL, and 4) all three simultaneously at a final concentration of 5 mg/mL. **(D)** Growth of pediatric and fetal organoids after apical stimulation with a combination of all three HMOs from day 0, compared to untreated organoids. The Sartorius SX5 Incucyte imaged each well every 6 hours and was used to quantify organoid growth as the total change in organoid area over five days (n = 3). **(E)** Forskolin-induced swelling of organoids (left) in the presence or absence of HMOs, represented as the total change in organoid area from the point of forskolin treatment (5 μmol/L) and normalised to media-treated organoids, measured using the Sartorius SX5 Incucyte (right). Three different pediatric and fetal organoid lines were used with n = 3-4 technical replicates per line. Data in (C-E) are mean ± SEM, for comparisons between two groups, Student’s T-tests were used; for more than two groups, analysis of variance (ANOVA) was used.

We evaluated the effects of three commercially available HMOs—2’-fucosyllactose (2’FL), 3’-sialyllactose (3’SL), and difucosyllactose (DFL)—using several functional assays (Fig. 1b). Basolateral stimulation of small intestinal organoids with various combinations of these HMOs indicated no adverse effects on organoid viability (Fig. 1c). Consequently, we proceeded with the combination therapy for all subsequent experiments.

*In vivo*, HMOs are delivered orally, contacting the apical membrane of the intestinal epithelium. To more accurately replicate this biological context, we initiated treatments immediately upon splitting, allowing the HMOs to interact with the apical membrane as the organoids formed closed lumen structures. We also used fetal-derived organoids to assess whether the effects of HMOs on IECs are specific to the developmental stage. This approach did not significantly impact organoid growth over a five-day period, regardless of developmental stage (Fig. 1d), indicating that apical HMO stimulation does not adversely affect growth. Additionally, we performed a forskolin-induced swelling assay to assess barrier integrity under both homeostatic and inflammatory conditions. To mimic inflammation, fully formed organoids were exposed to physiologically relevant levels of interferon-gamma (IFN-γ) from day 5. The observed swelling indicated that membrane integrity was preserved following HMO treatment across both fetal and pediatric organoids (Fig. 1e, Supplementary Fig. 1a). We further quantified the expression of several markers of organoid health under normal and inflammatory conditions. IL-8 was used as a marker of inflammation, BAX was used to assess apoptotic stress, ZO-1 was used to evaluate barrier integrity, and Ki67 was used as a proxy for proliferation (Supplementary Fig. 1b-e). HMO treatment reduced baseline BAX expression compared to untreated organoids and attenuated the increase in BAX expression following IFN-γ exposure. These results suggest that HMOs in such a blend may reduce cell death responses in the terminal ileum under inflammatory conditions, although the expression of a single marker cannot provide a comprehensive mechanistic understanding.

Overall, our findings indicate that HMOs are safe and well-tolerated in both pediatric and fetal organoids. We provide evidence that this combination of HMOs supports organoid viability and maintains, or in some cases enhances, intestinal epithelial barrier function under both steady-state and inflammatory conditions.

### HMOs Induce Developmental Stage-Specific Transcriptional Responses in Human Fetal and Pediatric IEO

Next, we sought to further investigate the impact of the HMO blend on the human intestinal epithelium by performing bulk RNA sequencing of fetal and pediatric IEOs co-cultured with the HMO blend for 7 days (Fig. 1a). Hierarchical clustering based on the 500 most variable genes effectively distinguished HMO-treated samples from untreated controls in both fetal and pediatric IEOs (Fig. 2a, 2b). Specifically, IEOs derived from each donor clustered separately based on their exposure to HMOs, in both fetal and pediatric epithelium. Principal component analysis (PCA) further confirmed clear separation between HMO-treated and untreated IEOs (Fig. 2c, 2d). These results suggest that, despite expected inter-patient variability, HMO exposure induces robust, developmental stage-specific transcriptomic changes.

**Figure 2.**
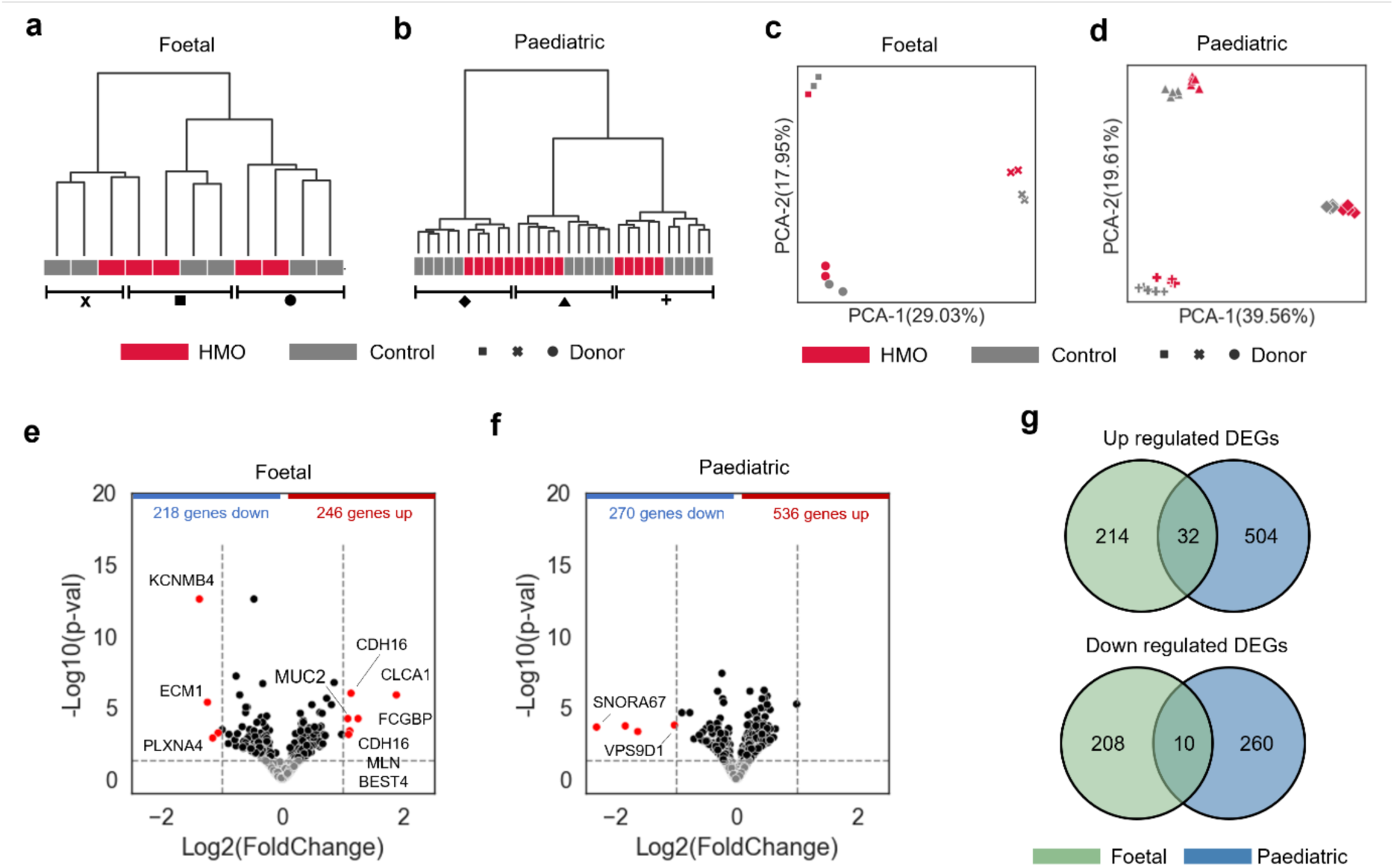
Transcriptomic differences between fetal IEOs and pediatric IEOs in response to HMO treatments. **(A, B)** Dendrograms representing hierarchical clustering of fetal IEO and pediatric IEO samples. Expression profiles of the top 500 genes with the highest variability in each group were used for the analysis. Colors indicate HMO treatment status, while symbols represent donors. **(C, D)** Scatter plots showing PCA results of fetal IEOs and pediatric IEO samples. Expression profiles of the top 500 genes with the highest variability in each group were used for the analysis. Colors indicate HMO treatment status, while symbols represent donors. **(E, F)** Volcano plots showing DEGs between HMO-treated and untreated samples in fetal IEOs and pediatric IEOs. The dotted line on the y-axis represents the threshold for DEGs (p-value < 0.05), while the two dotted lines on the x-axis represent the threshold for hDEGs (abs(Log_2_Fold-Change) > 1). Black dots represent DEGs, red dots indicate hDEGs, and grey dots represent all other genes. Gene names were labelled for coding genes among the hDEGs. **(G)** Venn diagram comparing DEGs in fetal IEOs and pediatric IEOs in response to HMO treatment. Upregulated and downregulated genes are shown separately.

To further elucidate the HMO-induced transcriptional changes at the individual gene level, we performed differential gene expression analysis. Of the 18,506 genes analyzed, HMO treatment significantly upregulated 246 genes and downregulated 218 genes in fetal IEOs. In contrast, in pediatric IEOs, 536 genes were upregulated and 270 downregulated compared to untreated controls (p-value < 0.05) (Fig. 2e, 2f, Supplementary Table 2). Among these, 10 genes in fetal IEOs and 4 genes in pediatric IEOs were highly differentially expressed genes (hDEGs), exhibiting substantial changes in expression (abs(Log_2_Fold-Change) > 1).

We then compared the differentially expressed genes (DEGs) between fetal and pediatric samples to assess age-related differences in the transcriptomic response to HMO treatment. Interestingly, the genes differentially expressed in response to the HMO blend showed significant variation between the two age groups. Of the 750 upregulated genes, only 32 (4.2%) were shared between fetal and pediatric organoids, and of the 478 downregulated genes, just 10 (2.1%) were common (Fig. 2g). Notably, none of the hDEGs were shared between the groups.

Overall, these findings imply that luminal exposure to HMOs can elicit distinct, age-specific transcriptional responses in the human intestinal epithelium.

### HMOs Can Induce Distinct Transcriptional Profiles Related to Lipid Metabolism in Pediatric IEOs

To further explore the impact of the HMO blend on pediatric and fetal IEOs at the pathway level, we performed an over-representation analysis (ORA) of DEGs using the Reactome database. Interestingly, no significant pathways were detected in fetal IEOs (Supplementary Table 3). In contrast, HMO treatment significantly affected pathways related to lipid metabolism and the transport of small molecules in pediatric organoids (p < 0.001) (Fig. 3a, Supplementary Table 4). These findings suggest that HMO-induced transcriptional changes in pediatric organoids predominantly affect metabolic processes, specifically those involving lipid handling.

**Figure 3.**
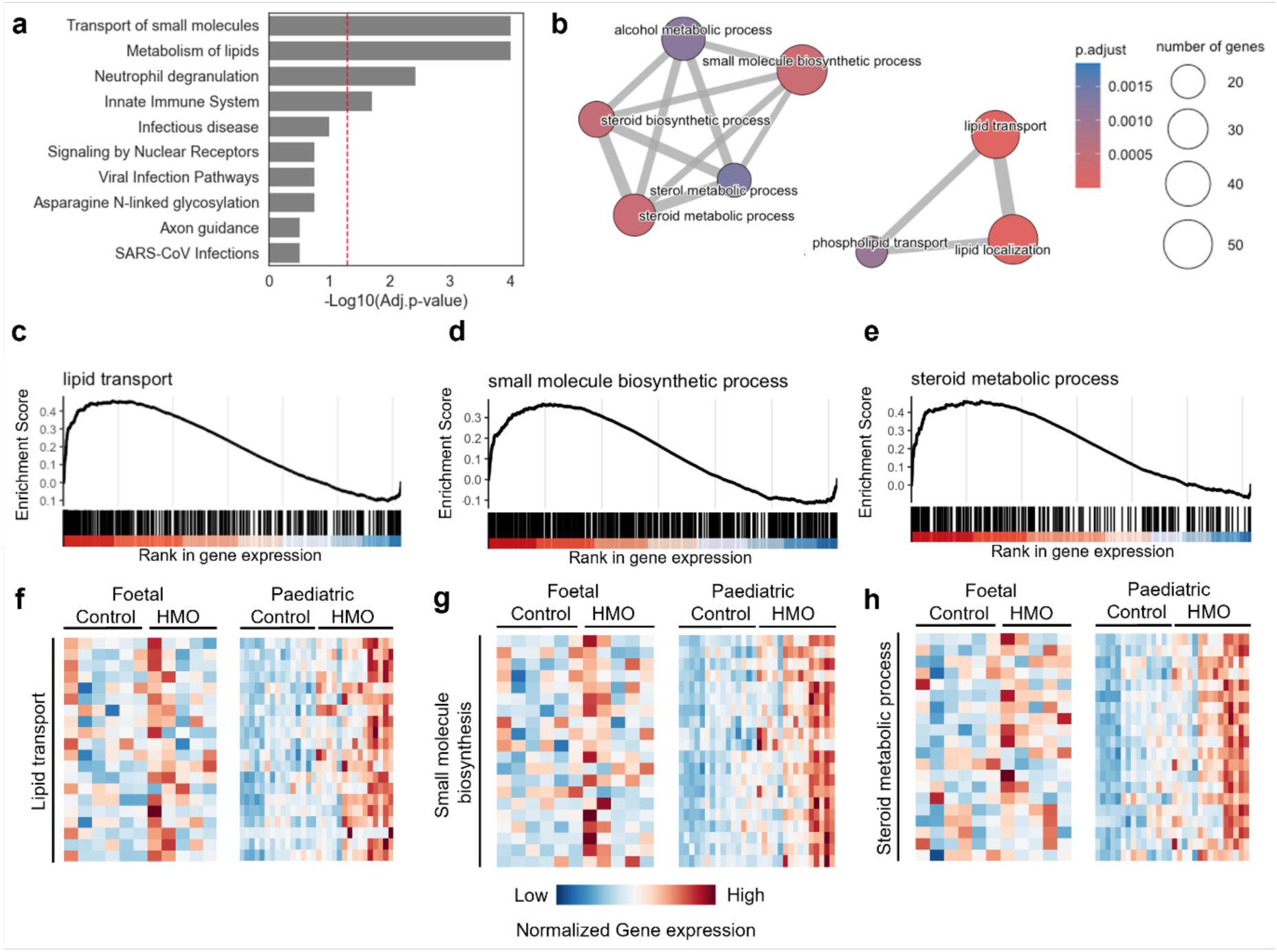
Biological pathways and processes affected by HMO treatment in pediatric IEOs. **(A)** Bar graph representing the top 10 pathways from the Reactome-based ORA results for pediatric IEOs. The red dashed line indicates the statistical significance threshold, with an adjusted p-value < 0.05. **(B)** Gene-sharing network visualizing the top 10 biological processes from the Gene Ontology-based ORA results for pediatric IEOs. Biological processes sharing more than 20% of genes are connected, and the edge thickness corresponds to the number of shared genes. Biological processes that did not form clusters are not displayed. **(C-E)** Gene Ontology-based GSEA results for pediatric IEOs. The top 3 overlapping biological processes from the ORA analysis are selectively shown. The color gradient at the bottom of each graph represents gene expression levels. **(F-H)** Heatmaps showing the expression levels of the top 20 genes most associated with each biological process for all IEO samples. Gene expression levels are normalized using z-scores.

To gain deeper insight into the biological processes modulated by HMOs in pediatric IEOs, we conducted a Gene Ontology-based ORA. This analysis revealed the top 10 significantly enriched biological processes (adjusted p-value < 0.05), which were organized into two densely connected network clusters (Fig. 3b, Supplementary Table 5). One cluster was associated with lipid transport and localization, while the other involved lipid metabolic processes. These findings mirror the top Reactome pathways, highlighting lipid-related biological activities as central to the HMO response in pediatric IEOs.

Given the prominent impact of the tested HMOs on lipid metabolism, we further employed Gene Set Enrichment Analysis (GSEA) to identify the directionality of gene expression changes within the affected processes. GSEA confirmed significant activation of key biological processes, including lipid transport (GO:0006869), steroid metabolic process (GO:0008202), and small molecule biosynthesis (GO:0044283) (Fig. 3c-e, Supplementary Table 6). A heatmap visualization of the top 20 genes involved in these processes demonstrated a consistent overexpression of lipid metabolism-related genes in HMO-treated pediatric IEOs (Fig. 3f-h). Notably, such transcriptional changes were not observed in fetal IEOs, reinforcing the idea that HMOs exert stage-specific effects.

Overall, these results demonstrate that the HMO blend most clearly induces lipid metabolism pathways in pediatric IEOs, with no comparable pathway effect observed in fetal IEOs.

### Developmental Stage-Specific Transcriptomic Changes to HMOs in Baseline and Inflammatory Conditions

We performed bulk RNA sequencing on fetal and pediatric IEOs treated with HMOs and IFN-γ to investigate the transcriptomic effects of the HMO blend under both baseline and inflammatory conditions. In this experimental setup, IEOs were first cultured with HMOs and then exposed to IFN-γ to simulate an inflammatory environment. PCA of all samples revealed that the IEOs effectively captured the transcriptomic effects of IFN-γ treatment (Fig. 4a). Additionally, hierarchical clustering and PCA of IFN-γ-treated samples showed that HMO treatment continued to alter the transcriptome in both fetal and pediatric IEOs within the inflammatory context (Fig. 4b, 4c, Supplementary Fig. 2a, 2b). Although the separation between HMO-treated and untreated samples was less pronounced under inflammation, the organoid model was still able to detect subtle HMO-induced transcriptomic changes despite the strong perturbation caused by IFN-γ exposure.

**Figure 4.**
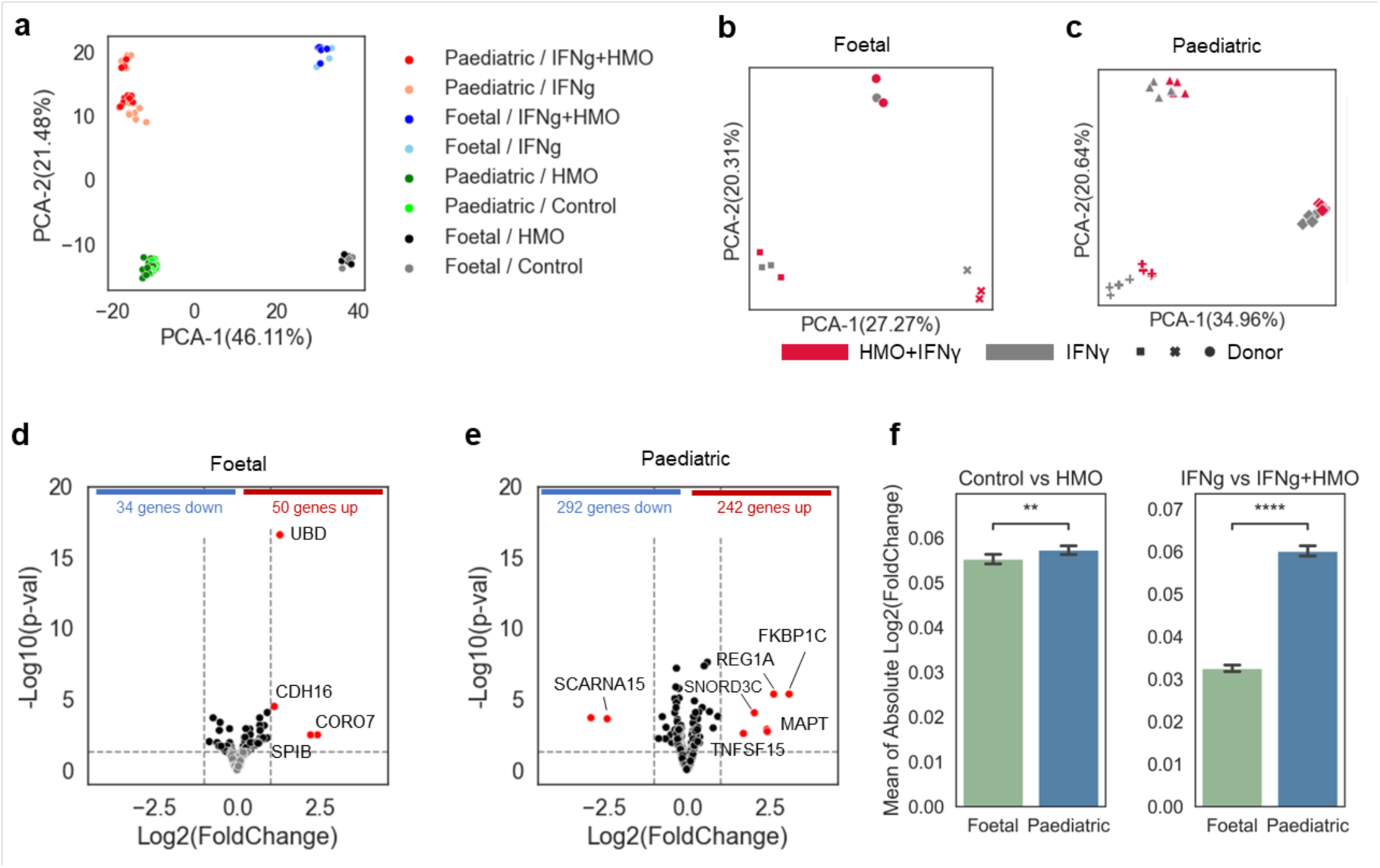
Transcriptomic differences between fetal IEOs and pediatric IEOs in response to HMO treatment under IFN-γ-induced inflammation. **(A)** Scatter plots showing the PCA results of all fetal IEO and pediatric IEO samples, with donor age and experimental conditions distinguished by colors. **(B, C)** Scatter plots showing PCA results of fetal IEOs and pediatric IEOs samples in the context of inflammation under IFN-γ induced inflammation. Expression profiles of the top 500 genes with the highest variability in each group were used for the analysis. Colors indicate HMO treatment status, while symbols represent donors. **(D, E)** Volcano plots showing DEGs between HMO-treated and untreated samples in fetal IEOs and pediatric IEOs under IFN-γ induced inflammation. The dotted line on the y-axis represents the threshold for DEGs (p-value < 0.05), while the two dotted lines on the x-axis represent the threshold for hDEGs (abs(Log_2_Fold-Change) > 1). Black dots represent DEGs, red dots indicate hDEGs, and grey dots represent all other genes. Gene names were labelled for coding genes among the hDEGs. **(F)** Bar graphs representing the differential effects of IFN-γ treatment on transcriptomic changes induced by HMOs in fetal IEOs and pediatric IEOs. A paired sample T-test was used to calculate statistical significance.

Differential gene expression analysis revealed significant transcriptional changes in both fetal and pediatric IEOs. In fetal IEOs, HMO treatment led to the upregulation of 50 genes and downregulation of 34 genes, while pediatric IEOs exhibited more pronounced changes, with 242 upregulated genes and 292 downregulated genes (Fig. 4d, 4e, Supplementary Table 2). Among these, 4 genes in fetal IEOs and 10 in pediatric IEOs displayed substantial changes in expression (abs(Log_2_Fold-Change) > 1). Once again, the transcriptional response to the HMO blend varied considerably between the two developmental stages. Only 4 upregulated and 2 downregulated genes were shared between fetal and pediatric IEOs (Supplementary Fig. 2c), highlighting distinct, stage-specific responses to the blend of HMOs.

Pediatric IEOs exhibited a robust and consistent response to the HMO blend, even under IFN-γ-induced inflammation. In contrast, fetal IEOs displayed fewer transcriptional changes overall. When we quantified the impact of HMO treatment on the transcriptome, pediatric IEOs maintained a stable transcriptional response regardless of IFN-γ treatment. However, in fetal IEOs, the influence of the HMO blend on transcriptomic changes was reduced in the presence of IFN-γ (p-value < 1.0×10^-4^), suggesting that HMO-modulation of gene expression in fetal IEOs is more susceptible to inflammatory disruption (Fig. 4f).

Taken together, these results demonstrate that the HMO blend can induce distinct, developmental stage-specific transcriptomic changes in both baseline and inflammatory conditions, with pediatric IEOs showing a more resilient and consistent response compared to fetal IEOs.

### HMOs Suppress Inflammation-Induced Activation of Protein Translation Pathways

To investigate the effects of HMOs on IEOs during inflammation, we performed Reactome-based over-representation analysis (ORA) of DEGs in both fetal and pediatric IEOs treated with the HMO blend and IFN-γ. While no pathways were found to be significantly enriched in fetal IEOs (Supplementary Table 7), pediatric IEOs displayed significant enrichment in pathways associated with cellular responses to stress, translation, and protein synthesis (Fig. 5a, Supplementary Table 8). Notably, the most significantly affected pathway in pediatric IEOs was the “Translation” pathway (R-HSA-72766), suggesting that the HMO blend may modulate the inflammatory response by regulating protein synthesis.

**Figure 5.**
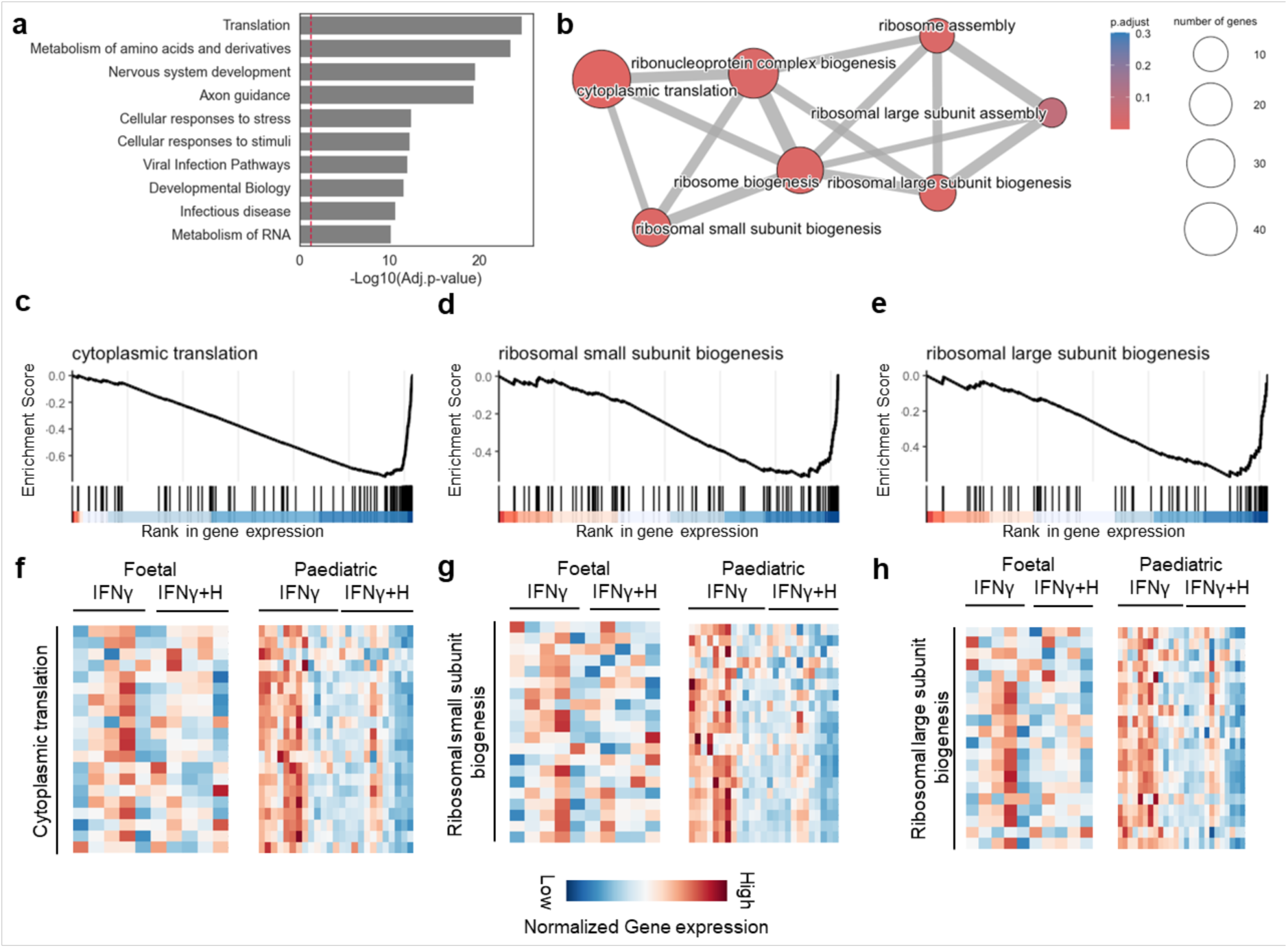
Biological pathways and processes affected by HMO treatment in pediatric IEOs under IFN-γ induced inflammation. **(A)** Bar graph representing the top 10 pathways from the Reactome-based ORA results for pediatric IEOs under inflammatory conditions by IFN-γ treatment. The red dashed line indicates the statistical significance threshold, with an adjusted p-value < 0.05. **(B)** Gene-sharing network visualizing the top 10 biological processes from the Gene Ontology-based ORA results for pediatric IEOs under inflammatory conditions. Biological processes sharing more than 20% of genes are connected, and the edge thickness corresponds to the number of shared genes. Biological processes that did not form clusters are not displayed. **(C-E)** Gene Ontology-based GSEA results for pediatric IEOs under inflammatory conditions. The top 3 overlapping biological processes from the ORA analysis are selectively shown. The color gradient at the bottom of each graph represents gene expression levels. **(F-H)** Heatmaps showing the expression levels of the top 20 genes most associated with each biological process for all IEO samples. Gene expression levels are normalized using z-scores.

Gene Ontology-based ORA further confirmed these findings, revealing enriched biological processes related to ribosome biogenesis, ribonucleoprotein complex assembly, and cytoplasmic translation (Fig. 5b, Supplementary Table 9). These processes are critical for protein synthesis, and their downregulation suggests that the HMO blend may suppress the excessive translation that is often induced during inflammation. GSEA validated these results, showing significant inhibition of biological processes involved in ribosomal function and protein synthesis in HMO-treated pediatric IEOs under inflammatory conditions (Fig. 5c-e, Supplementary Table 10).

A heatmap of key genes involved in translation processes revealed consistent downregulation in pediatric IEOs pre-treated with HMOs (Fig. 5f-h). This suppression of translation-related gene expression may represent a protective mechanism through which HMOs mitigate the heightened protein synthesis and cellular stress responses typically triggered by inflammation. Interestingly, this effect was not observed in fetal IEOs, reinforcing the developmental stage-specific impact of HMOs.

Collectively, these results demonstrate that these HMOs can suppress the inflammation-induced activation of protein translation pathways, particularly in pediatric IEOs, potentially limiting excessive protein synthesis and mitigating cellular stress during inflammatory responses.

### HMOs Counteract IFN-γ-Induced Inflammatory Gene Expression in Pediatric IEOs

To specifically examine the impact of HMO pre-treatment on IFN-γ-induced inflammation in pediatric IEOs, we analyzed the overlap between DEGs affected by both IFN-γ treatment and HMO pre-treatment (Fig. 6a). In this analysis, we identified genes whose expression changes induced by IFN-γ were either enhanced or diminished by HMO pre-treatment. Of the 166 overlapping genes, the HMO blend mitigated the IFN-γ-induced expression changes in 110 genes (66.3%) (Fig. 6b, 6c). Specifically, 40 genes that were upregulated by IFN-γ were downregulated with HMOs, while 70 genes that were downregulated by IFN-γ were upregulated after pre-treatment with HMOs. For all six hDEGs (abs(Log_2_Fold-Change) > 0.5), HMOs reversed the expression changes induced by IFN-γ (Fig. 6c). These findings suggest that HMOs can significantly counteract the transcriptomic alterations caused by IFN-γ, indicating a potential preventive role of HMOs against inflammation.

**Figure 6.**
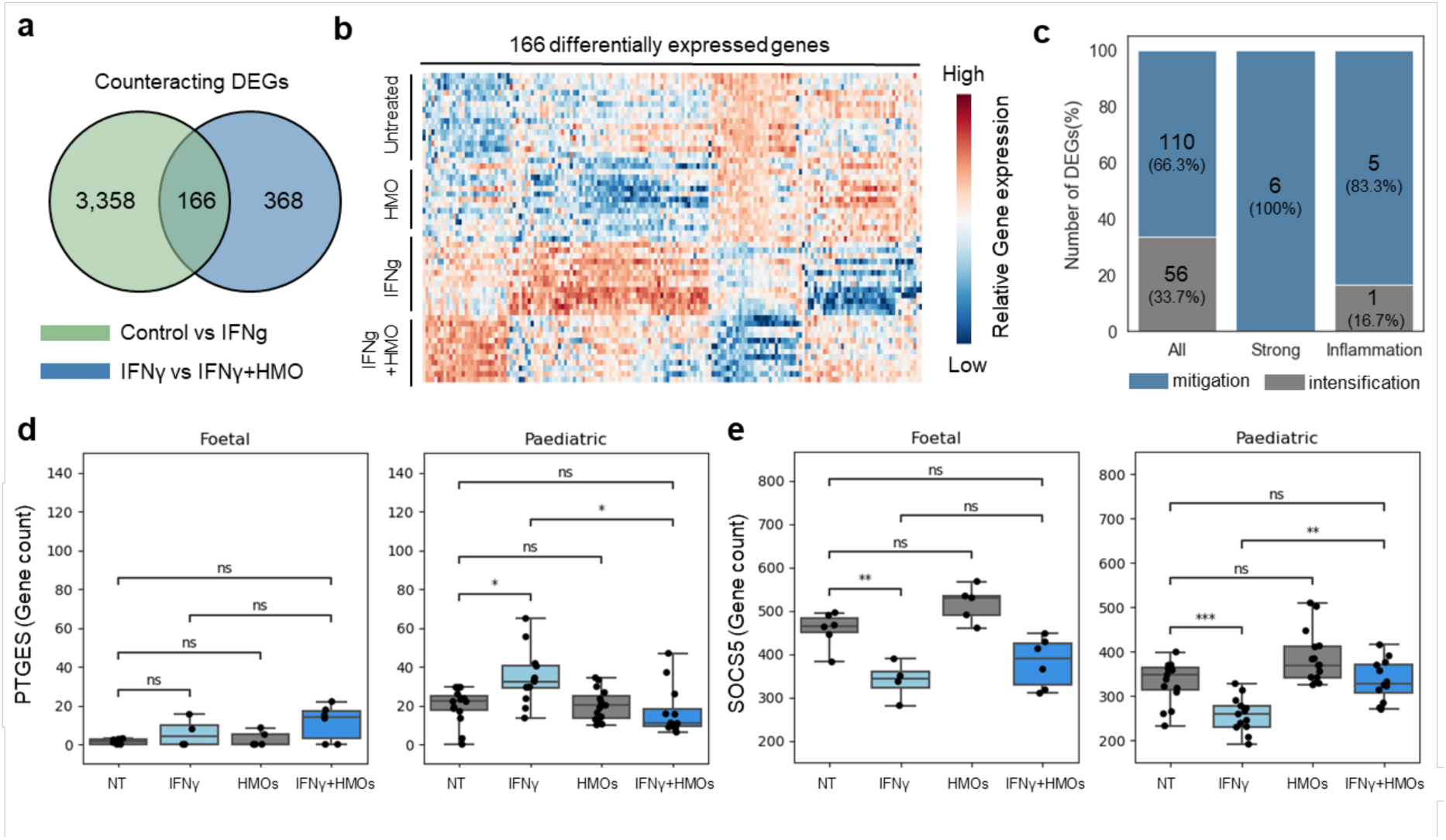
Genes with reduced IFN-γ-induced changes following HMO pre-treatment in pediatric IEOs. **(A)** Overlapping gene count between two DEG groups, with one group consisting of DEGs from IFN-γ treatment and the other from HMO treatment under IFN-γ-induced inflammation. **(B)** Heatmap showing the expression of the 166 overlapping genes identified in (A). The gene expression levels were Z-normalized, and the genes were ordered based on clustering according to their expression patterns. **(C)** Bar graph categorizing the 166 overlapping genes from (A) based on the direction of influence by HMOs. Genes, where HMO pre-treatment mitigated IFN-γ-induced changes, are represented in blue color, while those that intensified IFN-γ-induced changes are shown in grey color. Sub-statistics for six hDEGs (abs(Log_2_Fold-Change) > 1) and six inflammation-related genes are shown as separate bars. **(D, E)** Box plots displaying the expression levels of the PTGES and SOCS5 genes across all sample types. Gene expression levels are based on normalized gene count values.

Further analysis of inflammation-related genes revealed that the HMO blend attenuated IFN-γ-induced expression changes in all six targeted genes (Fig. 6c, Supplementary Table 11). Among these, the *PTGES* gene, which was upregulated by IFN-γ, showed a significant reduction (p-value < 0.05) in expression following HMO pre-treatment (Fig. 6d). This effect was specific to pediatric IEOs and was not observed in fetal IEOs. *PTGES* encodes the terminal enzyme in the cyclooxygenase (COX)-2-mediated prostaglandin E2 biosynthesis pathway, which is closely associated with intestinal inflammation and diseases such as ulcerative colitis [33]. Additionally, *SOCS5*, which was downregulated by IFN-γ, was significantly upregulated by HMOs in pediatric IEOs (p-value < 0.001). *SOCS5* is a member of the suppressor of cytokine signalling (SOCS) family, known to regulate immune responses and inflammation [34]. Taken together, these findings demonstrate that HMOs effectively counteract IFN-γ-induced inflammatory gene expression changes in pediatric IEOs, highlighting their potential as preventive agents against inflammation-driven diseases.

## DISCUSSION

Human milk oligosaccharides (HMOs) are known to play diverse physiological roles, including supporting a balanced infant gut microbiota, enhancing gastrointestinal barrier integrity, preventing infections, and potentially contributing to immune, brain, and cognitive development [35–39]. However, most prior studies have relied on animal models or cell lines, thereby limiting their relevance to human biology. In this study, we present novel findings on the effects of an HMO blend using IEOs derived from human fetal and pediatric tissues. Organoid models offer a highly relevant platform for HMO research, as they closely mimic human intestinal physiology and developmental stage, allowing for a more precise investigation of HMO-driven effects on gut development and immune responses [40]. These models provide ethical advantages over animal studies and enable personalized research using patient-derived cells [41], making them a valuable tool for exploring nutritional and therapeutic interventions aimed at improving gut development and health.

Our findings indicate that HMOs are not only safe and well-tolerated by fetal and pediatric IEOs under both steady-state and inflammatory conditions but also show potential in enhancing gut health and modulating inflammation pre- and post-natally. By utilizing these IEOs, we have revealed both age-specific and context-dependent effects of an HMO blend on intestinal epithelial cells. In pediatric IEOs, HMOs notably activated lipid metabolic pathways, aligning with the high lipid content of breast milk [42–43], suggesting that such HMOs may optimize lipid digestion and absorption and support efficient energy utilization during early childhood growth. In contrast, fetal IEOs displayed a more diffuse response to the HMO blend, underscoring developmental stage-specific differences in HMO activity. Nevertheless, the HMO blend in fetal IEOs produced gene signatures consistent with canonical goblet cell differentiation and mucus protein production, including Muc2, Fcgbp and Clca1, indicating a role in supporting epithelial cell differentiation and protecting against inflammation and infection [44–45]. In the context of inflammation, HMOs exhibited protective effects by attenuating the expression of key inflammation-related genes, including *PTGES*, which is linked to prostaglandin E2 biosynthesis and intestinal inflammation. HMOs also modulated translation-related pathways, suggesting a role in mitigating cellular stress and limiting tissue damage during inflammation [46–47]. These results emphasize the potential of HMOs as preventative agents against inflammation-related diseases, such as ulcerative colitis and Crohn’s Disease, by modulating inflammatory pathways and maintaining intestinal homeostasis.

Our study’s findings also hold significant implications for the development of breast milk substitutes and nutritional supplements. The age-specific effects of the tested HMOs point to the possibility of nutrition strategies that could be tailored to different developmental stages, optimizing both gut health and immune support in infants. By enhancing lipid metabolism and counteracting inflammatory responses, HMOs represent a promising approach to comprehensive early-life nutrition.

While our study provides valuable insights, there are limitations to consider. IEOs, though more representative of human biology than 2D cell lines, cannot fully replicate the complexity of the *in vivo* human gut. Further research could benefit from incorporating more diverse IEO models, including those from various genetic backgrounds, to better understand the mechanistic pathways influenced by HMOs. Additionally, clinical studies are needed to validate these findings in real-world settings and to assess the long-term impact of HMO supplementation on infant growth and health outcomes.

In conclusion, our study highlights the potential of HMOs to support gut health, with distinct effects observed across different developmental stages. By using human IEO models, we have shown that HMOs can enhance lipid metabolism and mitigate inflammatory responses, underscoring their value as key components of infant nutrition including preterm infant nutrition. As research into the functionality of HMOs advances, these insights could inform the development of innovative nutritional strategies that promote health and prevent disease from the earliest stages of life.

## METHODS

### PATIENT RECRUITMENT AND SAMPLE COLLECTION

Patients from the pediatric gastroenterology unit at Addenbrooke’s Hospital consented to provide additional biopsies from the terminal ileum (TI) with full ethical approval (REC-12/EE/0482). Children with macroscopically and histologically normal mucosa were selected. Proximal (small intestine) fetal gut was obtained with ethical approval (REC-96/085) and informed consent from elective terminations at 8-12 weeks gestational age. Sample and patient information are provided in Supplementary Table 1.

### HUMAN INTESTINAL EPITHELIAL ORGANOID CULTURE

Human intestinal epithelial organoids (IEOs) were generated from mucosal biopsy specimens and cultured in Matrigel (Corning) according to published protocols [32, 48–50]

### ORGANOID CO-CULTURE WITH HUMAN MILK OLIGOSACCHARIDES AND PRO-INFLAMMATORY CYTOKINES

Human milk oligosaccharides were kindly provided by dsm-firmenich, Switzerland. 500 mg/mL stocks of 2’-fucosyllactose (2’FL), 3’-sialyllactose (3’SL) and 2’-fucosyllactose/difucosyllactose (2’FL/DFL) in PBS were diluted to a final concentration of 5 mg/mL in organoid complete medium.

Human IEOs were cultured for five days and supplemented with human milk oligosaccharides every other day to contact the apical surface of the epithelium. To explore the effect of inflammation, IEOs were treated with IFN-γ (PHC4031; Life Technologies, Carlsbad, CA) at 20 ng/mL for the final 48 hours.

### HUMAN IEO GROWTH AND BARRIER INTEGRITY ASSESSMENT

IEO growth in the presence or absence of HMOs was imaged using the Incucyte SX5 (Sartorius AG, Göttingen, Germany) every 6 hours for 7 days. The change in organoid area relative to the first image was calculated using the Incucyte IEO analysis software to quantify organoid growth over time. At least three wells were analysed to generate an average measurement of IEO growth for each line.

Organoids were treated with 5 μmol/L forskolin on day 7 to induce swelling and assess the barrier integrity of the HMO-treated organoids. The organoids were imaged every 4 hours for 48 hours in the Incucyte SX5 and the swelling was quantified as above.

### HUMAN IEO VIABILITY

Organoids were seeded in 96 well plates and exposed to various HMO combinations at a final concentration of 5 mg/mL. On day 8, the viability was quantified using the CellTiter-Glo3D assay. This assay generates a luminescent readout based on the amount of ATP present as a marker of metabolically active cells. The experiment included three repeats per condition and empty Matrigel domes as a negative control.

### HARVESTING HUMAN IEOS AND RNA EXTRACTION

At the end of each experiment, organoids were harvested and stored in AllProtect tissue reagent at −80°C. RNA was isolated using the GenElute Mammalian Total RNA Miniprep Kit (Sigma). Samples for bulk RNA sequencing included an additional on-column DNase I digestion step (Sigma).

### GENE EXPRESSION ANALYSIS USING QUANTITATIVE PCR

cDNA was prepared using Superscript III Reverse Transcriptase (Invitrogen) and used for quantitative PCR using Quantifast SYBR Green (Qiagen). The mRNA expression of each gene was normalised to the expression of a housekeeping gene, GAPDH, and relative expression was analysed using the ΔΔCt method [51]. Primer sequences are described in Table 1.

**Table 1.**
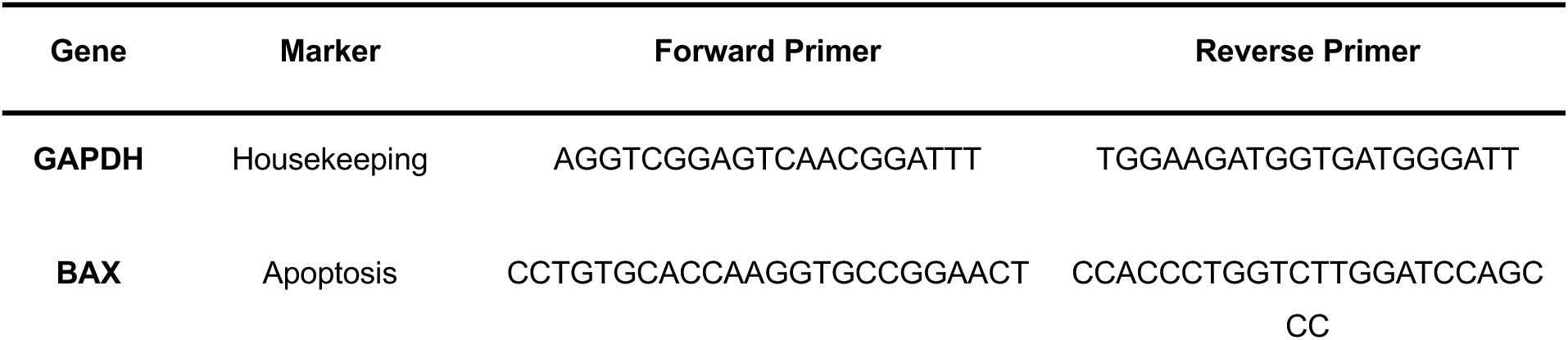
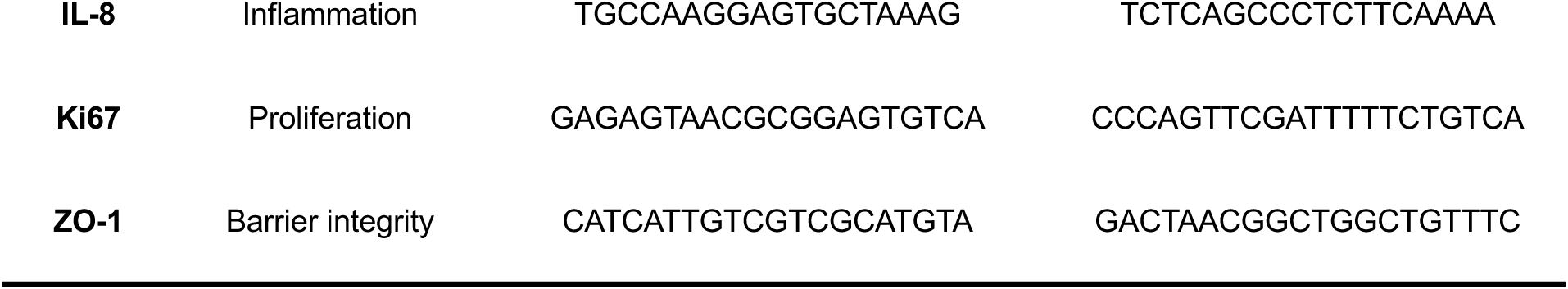
Primers used for quantitative qPCR of samples.

### BULK RNA SEQUENCING

RNA sequencing was performed by Cambridge Genomic Services, University of Cambridge. The libraries were sequenced on a NextSeq 2000 platform using a P3 50 cycle sequencing kit. The sequencing was single end with a read length of 50 bases.

### RNA SEQUENCING DATA PREPROCESSING

RNA sequencing data was processed using nf-core/rnaseq v3.14.0 (doi: https://doi.org/10.5281/zenodo.1400710) from the nf-core collection of workflows [52], utilizing reproducible software environments from the Bioconda [53] and Biocontainers [54] projects. The pipeline was executed with Nextflow v23.10.1 [55]. We used GRCh37 as the reference genome, STAR [56] for alignment, and Salmon [57] for quantification. Other parameters were set to the default values suggested in the nf-core/rnaseq pipeline.

### DIFFERENTIALLY EXPRESSED GENE ANALYSIS USING BULK RNA SEQUENCING DATA

Differentially expressed gene (DEG) analysis was performed using the nf-core/differentialabundance v1.40 pipeline (doi: 10.5281/zenodo.7568000) from the nf-core collection of workflows. The pipeline was executed with Nextflow v23.10.1 [55]. Genes were excluded from the analysis if their expression (count) was less than 1 in more than 50% of samples (min_proportion). Outlier samples detected during quality control with an absolute median absolute deviation score > 5 were excluded from further analysis. DEGs were identified based on an adjusted p-value < 0.05, and those with abs(Log_2_Fold-Change) > 1 were classified as highly differentially expressed genes (hDEGs). All other parameters were set to default values recommended in the pipeline.

### SAMPLE CLUSTERING, OVER-REPRESENTATION ANALYSIS AND GENE SET ENRICHMENT ANALYSIS

The Python package Scikit-learn v1.4.1 [58] was utilized for PCA and hierarchical clustering. The top 500 genes with the highest variance between samples were selectively used for both analyses. The R package clusterProfiler v4.10.1 [59] was employed for ORA and GSEA. For Reactome-based ORA, pathways containing between 200 and 1,000 genes were considered. For Gene Ontology-based ORA and GSEA, biological processes containing between 10 and 500 genes were included. In ORA and GSEA, the threshold for detecting significantly related gene sets was set at an adjusted p-value < 0.05.

### STATISTICS AND REPRODUCIBILITY

Statistical analysis was performed using GraphPad Prism, Python or R. Plots containing confidence intervals represent mean ± SEM, unless indicated otherwise. For comparisons between two groups, Student’s T-tests were used; for more than two groups, analysis of variance (ANOVA) was used. Under these conditions, the assumption of normality was confirmed using the Shapiro-Wilk test. In cases where multiple comparisons were conducted, the adjusted p-value was used to control the false discovery rate. For heatmap visualization of gene expression, Z-Score normalization was applied to the expression abundance of each gene. To ensure reproducibility across all analyses, the random seed was fixed at 777.

### DATA VISUALIAZATION

The visualization of GSEA results was performed using the R package Enrichplot v1.22.0 [60], while other plots were generated using the Python package Seaborn v0.11.2 [61].

## ACKNOWLEDGMENTS

Processing of human tissues and generation of human IEOs was supported by the Helmsley Charitable Trust, funded by The Leona M and Harry B Helmsley Charitable Trust grant (G118500). All the experiments are funded by dsm-firmenich (G116308). JP and KN are funded by The Leona M and Harry B Helmsley Charitable Trust grant (G118500). ES is funded by the Cystic Fibrosis Trust in collaboration with the University of Sheffield (G107734). TD is funded by the Milner Therapeutics Institute.

## AUTHOR CONTRIBUTIONS

E.S., K.N. and T.D. performed the experiments and contributed to data curation. J.P. performed the data analysis and handled data curation. E.S., J.P. and K.N. contributed to the writing of the manuscript. M.Z. designed and led the study, secured funding, led data analyses and experimental work, and contributed to revising the final version of the manuscript. All authors approved the final version of the manuscript.

## COMPETING INTERESTS

The authors declare no competing interests.

## CORRESPONDANCE

Correspondence to Prof. Matthias Zilbauer.

## Notes

### Competing Interest Statement

The authors have declared no competing interest.

