## Supplementary Figures for "Human Milk Oligosaccharides Modulate Inflammatory Responses and Lipid Metabolism in a Human Intestinal Organoid Model"

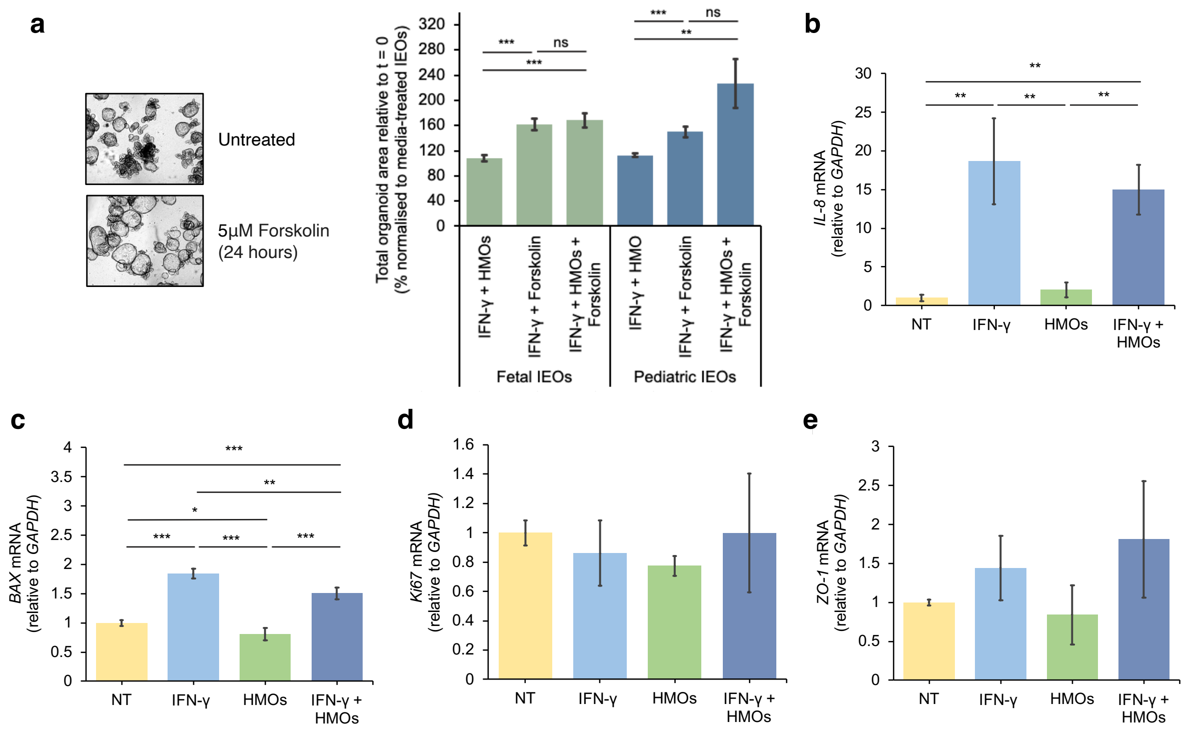


**Supplementary Figure 1.** Human milk oligosaccharides (HMOs) support organoid viability and maintain, and in some cases improve, cellular function under steady-state and inflammatory conditions.

**(A)** Forskolin-induced swelling of inflamed organoids in the presence or absence of HMOs, represented as the total change in organoid area from the point of forskolin treatment (5 µmol/L) and normalised to media-treated organoids, measured using the Sartorius SX5 Incucyte. To mimic inflammation, organoids were exposed to 20 ng/mL IFN-γ from day 5. Three different pediatric and fetal organoid lines were used with n = 3-4 technical replicates per line.

**(B-E)** *IL-8* (Supplementary Fig. 1b), *BAX* (Supplementary Fig. 1c), *Ki67* (Supplementary Fig. 1d) and *ZO-1* (Supplementary Fig. 1e) mRNA expression in the presence or absence of HMOs during steady-state or inflammation (n = 3). Organoids were treated with 20 ng/mL IFN-γ at day 5 to mimic inflammation.

Data in (B-E) are mean ± SEM, for comparisons between two groups, Student's T-tests were used; for more than two groups, analysis of variance (ANOVA) was used.

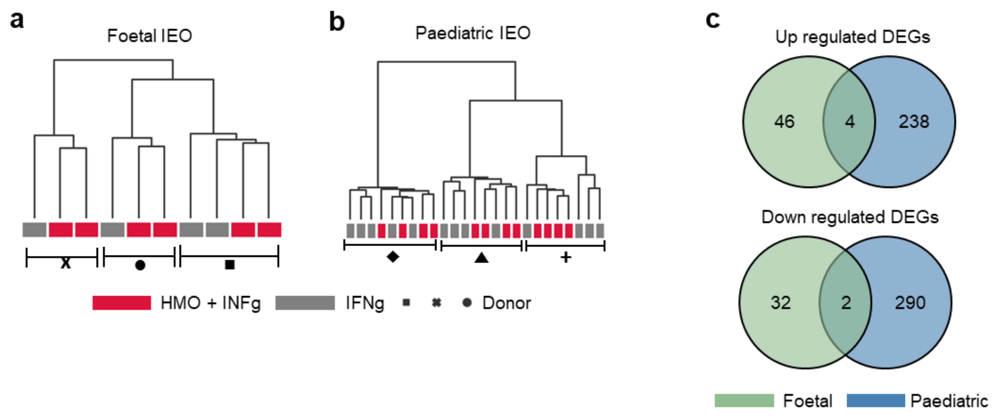


**Supplementary Figure 2.** Transcriptomic differences between fetal IEOs and pediatric IEOs in response to HMO treatment under IFN-γ-induced inflammation.

**(A, B)** Dendrograms representing hierarchical clustering of fetal IEO and pediatric IEO samples. Expression profiles of the top 500 genes with the highest variability in each group were used for the analysis. Colors indicate HMO treatment status, while symbols represent donors.

**(C)** Venn diagram comparing DEGs in fetal IEOs and pediatric IEOs in response to HMO treatment under IFN-γ induced inflammation. Upregulated and downregulated genes are shown separately.
