## Supplementary Tables for "Human Milk Oligosaccharides Modulate Inflammatory Responses and Lipid Metabolism in a Human Intestinal Organoid Model"

**Supplementary Table 1.** Sample information for organoids submitted for bulk RNA sequencing.

***PEDIATRIC SAMPLES***

| Sample | Age | Sex | Diagnosis | Gut Segment | Passage |
|---|---|---|---|---|---|
| 1 | 6 | Female | Control | Terminal ileum | 8 |
| 2 | 6 | Male | Control | Terminal ileum | 7 |
| 3 | 6 | Male | Control | Terminal ileum | 4 |

***FETAL SAMPLES***

| Sample | Gestational Age | Diagnosis | Segment | Passage |
|---|---|---|---|---|
| 1 | 11 weeks 3 days | Control | Ileum | 5 |
| 2 | 10 weeks 5 days | Control | Ileum | 5 |
| 3 | 10 weeks 4 days | Control | Ileum | 5 |

**Supplementary Table 2.** Statistics of differentially expressed genes (p-value < 0.05).

| Age | Condition | hDEGs* | Up regulated genes | Down regulated genes | All |
|---|---|---|---|---|---|
| Fetal | NT vs HMOs | 10 | 246 | 218 | 464 |
| Fetal | NT vs IFN-γ | 575 | 1,520 | 1,204 | 2,724 |
| Fetal | NT vs IFN-γ+HMOs | 688 | 2,180 | 2,056 | 4,236 |
| Fetal | IFN-γ vs IFN-γ+HMOs | 4 | 50 | 34 | 84 |
| Pediatric | NT vs HMOs | 4 | 536 | 270 | 806 |
| Pediatric | NT vs IFN-γ | 505 | 1,819 | 1,705 | 3,524 |
| Pediatric | NT vs IFN-γ+HMOs | 523 | 2,276 | 2,155 | 4,431 |
| Pediatric | IFN-γ vs IFN-γ+HMOs | 10 | 242 | 292 | 534 |

* hDEGs: Highly differentially expressed genes (abs($Log_2$Fold-Change) > 1 and p-value < 0.05)

* NT: Non-treated samples as control

**Supplementary Table 3.** Reactome-based ORA result (Fetal, NT vs HMOs).

| Rank | ID | Description | Adjusted p-value |
|---|---|---|---|
| 1 | R-HSA-168249 | Innate Immune System | 0.082922 |
| 2 | R-HSA-6798695 | Neutrophil degranulation | 0.082922 |
| 3 | R-HSA-72163 | mRNA Splicing - Major Pathway | 0.082922 |
| 4 | R-HSA-72172 | mRNA Splicing | 0.088783 |
| 5 | R-HSA-112316 | Neuronal System | 0.139279 |
| 6 | R-HSA-1266738 | Developmental Biology | 0.160095 |
| 7 | R-HSA-9675108 | Nervous system development | 0.165272 |
| 8 | R-HSA-72203 | Processing of Capped Intron-Containing Pre-mRNA | 0.221057 |
| 9 | R-HSA-422475 | Axon guidance | 0.241236 |
| 10 | R-HSA-9006931 | Signaling by Nuclear Receptors | 0.276132 |
| 11 | R-HSA-5653656 | Vesicle-mediated transport | 0.276132 |
| 12 | R-HSA-5663202 | Diseases of signal transduction by growth factor receptors and second messengers | 0.296236 |
| 13 | R-HSA-9006934 | Signaling by Receptor Tyrosine Kinases | 0.313575 |
| 14 | R-HSA-2262752 | Cellular responses to stress | 0.315154 |
| 15 | R-HSA-199991 | Membrane Trafficking | 0.315154 |
| 16 | R-HSA-8953854 | Metabolism of RNA | 0.315154 |
| 17 | R-HSA-8953897 | Cellular responses to stimuli | 0.327241 |
| 18 | R-HSA-556833 | Metabolism of lipids | 0.338048 |
| 19 | R-HSA-5683057 | MAPK family signaling cascades | 0.342973 |
| 20 | R-HSA-5663205 | Infectious disease | 0.354336 |
| 21 | R-HSA-68882 | Mitotic Anaphase | 0.390682 |
| 22 | R-HSA-2555396 | Mitotic Metaphase and Anaphase | 0.390682 |
| 23 | R-HSA-73894 | DNA Repair | 0.390682 |
| 24 | R-HSA-9824446 | Viral Infection Pathways | 0.405326 |
| 25 | R-HSA-71387 | Metabolism of carbohydrates | 0.405326 |
| 26 | R-HSA-9006925 | Intracellular signaling by second messengers | 0.405326 |
| 27 | R-HSA-382551 | Transport of small molecules | 0.405326 |
| 28 | R-HSA-388396 | GPCR downstream signalling | 0.405326 |
| 29 | R-HSA-449147 | Signaling by Interleukins | 0.405326 |
| 30 | R-HSA-162906 | HIV Infection | 0.405326 |

**Supplementary Table 4.** Reactome-based ORA result (Pediatric, NT vs HMOs).

| Rank | ID | Description | Adjusted p-value |
|---|---|---|---|
| 1 | R-HSA-382551 | Transport of small molecules | 0.000097 |
| 2 | R-HSA-556833 | Metabolism of lipids | 0.000097 |
| 3 | R-HSA-6798695 | Neutrophil degranulation | 0.00376 |
| 4 | R-HSA-168249 | Innate Immune System | 0.019557 |
| 5 | R-HSA-5663205 | Infectious disease | 0.098357 |
| 6 | R-HSA-9006931 | Signaling by Nuclear Receptors | 0.171676 |
| 7 | R-HSA-9824446 | Viral Infection Pathways | 0.171676 |
| 8 | R-HSA-446203 | Asparagine N-linked glycosylation | 0.176394 |
| 9 | R-HSA-422475 | Axon guidance | 0.305225 |
| 10 | R-HSA-9679506 | SARS-CoV Infections | 0.305225 |
| 11 | R-HSA-9012999 | RHO GTPase cycle | 0.305225 |
| 12 | R-HSA-9675108 | Nervous system development | 0.386138 |
| 13 | R-HSA-9716542 | Signaling by Rho GTPases, Miro GTPases and RHOBTB3 | 0.386138 |
| 14 | R-HSA-194315 | Signaling by Rho GTPases | 0.386138 |
| 15 | R-HSA-109582 | Hemostasis | 0.742301 |
| 16 | R-HSA-71291 | Metabolism of amino acids and derivatives | 0.742301 |
| 17 | R-HSA-72766 | Translation | 0.742301 |
| 18 | R-HSA-9694516 | SARS-CoV-2 Infection | 0.789641 |
| 19 | R-HSA-1266738 | Developmental Biology | 0.82267 |
| 20 | R-HSA-372790 | Signaling by GPCR | 0.898954 |
| 21 | R-HSA-8953897 | Cellular responses to stimuli | 0.898954 |
| 22 | R-HSA-2262752 | Cellular responses to stress | 0.928906 |
| 23 | R-HSA-5653656 | Vesicle-mediated transport | 0.951171 |
| 24 | R-HSA-71387 | Metabolism of carbohydrates | 0.968056 |
| 25 | R-HSA-199991 | Membrane Trafficking | 0.980888 |
| 26 | R-HSA-388396 | GPCR downstream signaling | 0.999974 |
| 27 | R-HSA-112316 | Neuronal System | 0.999974 |
| 28 | R-HSA-913531 | Interferon Signaling | 0.999974 |
| 29 | R-HSA-8953854 | Metabolism of RNA | 0.999974 |
| 30 | R-HSA-195258 | RHO GTPase Effectors | 0.999974 |

**Supplementary Table 5.** Gene Ontology-based ORA result (Pediatric, NT vs HMOs).

| Rank | ID | Description | Adjusted p-value |
|---|---|---|---|
| 1 | GO:0010876 | lipid localization | 0.000002 |
| 2 | GO:0006869 | lipid transport | 0.000002 |
| 3 | GO:0008202 | steroid metabolic process | 0.000363 |
| 4 | GO:0044283 | small molecule biosynthetic process | 0.000363 |
| 5 | GO:0006694 | steroid biosynthetic process | 0.000419 |
| 6 | GO:0015914 | phospholipid transport | 0.001052 |
| 7 | GO:0006066 | alcohol metabolic process | 0.001223 |
| 8 | GO:0016125 | sterol metabolic process | 0.00136 |
| 9 | GO:0050892 | intestinal absorption | 0.001756 |
| 10 | GO:0006879 | intracellular iron ion homeostasis | 0.001839 |
| 11 | GO:1901615 | organic hydroxy compound metabolic process | 0.002336 |
| 12 | GO:0016126 | sterol biosynthetic process | 0.002336 |
| 13 | GO:0042632 | cholesterol homeostasis | 0.002817 |
| 14 | GO:1902652 | secondary alcohol metabolic process | 0.002817 |
| 15 | GO:0055092 | sterol homeostasis | 0.002817 |
| 16 | GO:0008203 | cholesterol metabolic process | 0.002817 |
| 17 | GO:0006641 | triglyceride metabolic process | 0.005756 |
| 18 | GO:0022600 | digestive system process | 0.005756 |
| 19 | GO:1901617 | organic hydroxy compound biosynthetic process | 0.006034 |
| 20 | GO:0140115 | export across plasma membrane | 0.006224 |
| 21 | GO:0007586 | digestion | 0.006277 |
| 22 | GO:0015748 | organophosphate ester transport | 0.006277 |
| 23 | GO:0046394 | carboxylic acid biosynthetic process | 0.00918 |
| 24 | GO:0006814 | sodium ion transport | 0.00918 |
| 25 | GO:0016053 | organic acid biosynthetic process | 0.00918 |
| 26 | GO:0034368 | protein-lipid complex remodeling | 0.009843 |
| 27 | GO:0034369 | plasma lipoprotein particle remodeling | 0.009843 |
| 28 | GO:0015718 | monocarboxylic acid transport | 0.010156 |
| 29 | GO:0071827 | plasma lipoprotein particle organization | 0.010156 |
| 30 | GO:0051235 | maintenance of location | 0.011005 |

**Supplementary Table 6.** Gene Ontology-based GSEA result (Pediatric, NT vs HMOs).

| Rank | ID | Description | Enrichment Score | NES | Adjusted p-value |
|---|---|---|---|---|---|
| 1 | GO:0022613 | ribonucleoprotein complex biogenesis | -0.461206 | -2.262853 | 5.79E-15 |
| 2 | GO:0002181 | cytoplasmic translation | -0.602478 | -2.56938 | 1.71E-11 |
| 3 | GO:0042254 | ribosome biogenesis | -0.465543 | -2.195212 | 5.31E-10 |
| 4 | GO:0015711 | organic anion transport | 0.477495 | 2.066771 | 2.55E-09 |
| 5 | GO:1901615 | organic hydroxy compound metabolic process | 0.441828 | 1.95884 | 2.55E-09 |
| 6 | GO:0006091 | generation of precursor metabolites and energy | 0.434887 | 1.934168 | 2.55E-09 |
| 7 | GO:0034470 | ncRNA processing | -0.40522 | -1.9639 | 1.12E-08 |
| 8 | GO:0006869 | lipid transport | 0.456149 | 1.984027 | 1.50E-08 |
| 9 | GO:0006631 | fatty acid metabolic process | 0.466453 | 2.018912 | 2.43E-08 |
| 10 | GO:0010876 | lipid localization | 0.443978 | 1.958366 | 2.43E-08 |
| 11 | GO:0016072 | rRNA metabolic process | -0.469053 | -2.160896 | 3.70E-08 |
| 12 | GO:0071826 | protein-RNA complex organization | -0.494916 | -2.210284 | 4.37E-08 |
| 13 | GO:0022618 | protein-RNA complex assembly | -0.500742 | -2.21987 | 4.63E-08 |
| 14 | GO:0006066 | alcohol metabolic process | 0.462064 | 1.988928 | 1.16E-07 |
| 15 | GO:0009259 | ribonucleotide metabolic process | 0.443094 | 1.923531 | 1.16E-07 |
| 16 | GO:0009150 | purine ribonucleotide metabolic process | 0.45193 | 1.952733 | 2.44E-07 |
| 17 | GO:0019693 | ribose phosphate metabolic process | 0.439042 | 1.910645 | 2.79E-07 |
| 18 | GO:0015849 | organic acid transport | 0.478482 | 2.019656 | 3.52E-07 |
| 19 | GO:0046942 | carboxylic acid transport | 0.476232 | 2.008256 | 4.44E-07 |
| 20 | GO:0005975 | carbohydrate metabolic process | 0.399315 | 1.783697 | 4.44E-07 |
| 21 | GO:0006364 | rRNA processing | -0.477308 | -2.160589 | 4.82E-07 |
| 22 | GO:0006163 | purine nucleotide metabolic process | 0.42379 | 1.870737 | 4.82E-07 |
| 23 | GO:0010256 | endomembrane system organization | 0.397637 | 1.781944 | 5.26E-07 |
| 24 | GO:0008202 | steroid metabolic process | 0.462885 | 1.948111 | 1.38E-06 |
| 25 | GO:0019646 | aerobic electron transport chain | 0.602709 | 2.222646 | 1.78E-06 |
| 26 | GO:0042274 | ribosomal small subunit biogenesis | -0.572848 | -2.281039 | 1.93E-06 |
| 27 | GO:0015718 | monocarboxylic acid transport | 0.558396 | 2.14902 | 2.28E-06 |
| 28 | GO:0006879 | intracellular iron ion homeostasis | 0.656745 | 2.269774 | 2.73E-06 |
| 29 | GO:0009117 | nucleotide metabolic process | 0.390411 | 1.735052 | 3.19E-06 |
| 30 | GO:0016042 | lipid catabolic process | 0.447525 | 1.902686 | 3.54E-06 |

**Supplementary Table 7.** Reactome-based ORA result (Fetal, IFN-γ vs IFN-γ + HMOs).

| Rank | ID | Description | Adjusted p-value |
|---|---|---|---|
| 1 | R-HSA-5683057 | MAPK family signaling cascades | 0.937078 |
| 2 | R-HSA-382551 | Transport of small molecules | 0.937078 |
| 3 | R-HSA-5673001 | RAF/MAP kinase cascade | 0.937078 |
| 4 | R-HSA-5684996 | MAPK1/MAPK3 signaling | 0.937078 |
| 5 | R-HSA-556833 | Metabolism of lipids | 0.937078 |
| 6 | R-HSA-6798695 | Neutrophil degranulation | 0.937078 |
| 7 | R-HSA-199991 | Membrane Trafficking | 0.937078 |
| 8 | R-HSA-5653656 | Vesicle-mediated transport | 0.937078 |
| 9 | R-HSA-372790 | Signaling by GPCR | 0.937078 |
| 10 | R-HSA-5688426 | Deubiquitination | 0.937078 |
| 11 | R-HSA-71387 | Metabolism of carbohydrates | 0.937078 |
| 12 | R-HSA-9012999 | RHO GTPase cycle | 0.937078 |
| 13 | R-HSA-109582 | Hemostasis | 0.937078 |
| 14 | R-HSA-1280218 | Adaptive Immune System | 0.937078 |
| 15 | R-HSA-168249 | Innate Immune System | 0.937078 |
| 16 | R-HSA-1266738 | Developmental Biology | 0.937078 |
| 17 | R-HSA-3700989 | Transcriptional Regulation by TP53 | 0.937078 |
| 18 | R-HSA-8953854 | Metabolism of RNA | 0.937078 |
| 19 | R-HSA-5663202 | Diseases of signal transduction by growth factor receptors and second messengers | 0.937078 |
| 20 | R-HSA-162906 | HIV Infection | 0.937078 |
| 21 | R-HSA-8951664 | Neddylation | 0.937078 |
| 22 | R-HSA-194315 | Signaling by Rho GTPases | 0.937078 |
| 23 | R-HSA-1257604 | PIP3 activates AKT signaling | 0.937078 |
| 24 | R-HSA-9006931 | Signaling by Nuclear Receptors | 0.937078 |
| 25 | R-HSA-913531 | Interferon Signaling | 0.937078 |
| 26 | R-HSA-9716542 | Signaling by Rho GTPases, Miro GTPases and RHOBTB3 | 0.937078 |
| 27 | R-HSA-422475 | Axon guidance | 0.937078 |
| 28 | R-HSA-112316 | Neuronal System | 0.937078 |
| 29 | R-HSA-9824446 | Viral Infection Pathways | 0.937078 |
| 30 | R-HSA-9006925 | Intracellular signaling by second messengers | 0.937078 |

**Supplementary Table 8.** Reactome-based ORA result (Pediatric, IFN-γ vs IFN-γ + HMOs).

| Rank | ID | Description | Adjusted p-value |
|---|---|---|---|
| 1 | R-HSA-72766 | Translation | 2.01E-25 |
| 2 | R-HSA-71291 | Metabolism of amino acids and derivatives | 3.28E-24 |
| 3 | R-HSA-9675108 | Nervous system development | 2.74E-20 |
| 4 | R-HSA-422475 | Axon guidance | 4.78E-20 |
| 5 | R-HSA-2262752 | Cellular responses to stress | 3.54E-13 |
| 6 | R-HSA-8953897 | Cellular responses to stimuli | 5.61E-13 |
| 7 | R-HSA-9824446 | Viral Infection Pathways | 9.22E-13 |
| 8 | R-HSA-1266738 | Developmental Biology | 2.52E-12 |
| 9 | R-HSA-5663205 | Infectious disease | 2.29E-11 |
| 10 | R-HSA-8953854 | Metabolism of RNA | 6.62E-11 |
| 11 | R-HSA-9694516 | SARS-CoV-2 Infection | 2.83E-04 |
| 12 | R-HSA-9679506 | SARS-CoV Infections | 5.77E-04 |
| 13 | R-HSA-5663202 | Diseases of signal transduction by growth factor receptors and second messengers | 1.27E-01 |
| 14 | R-HSA-449147 | Signaling by Interleukins | 1.45E-01 |
| 15 | R-HSA-556833 | Metabolism of lipids | 2.25E-01 |
| 16 | R-HSA-1280215 | Cytokine Signaling in Immune system | 5.12E-01 |
| 17 | R-HSA-168249 | Innate Immune System | 6.11E-01 |
| 18 | R-HSA-195258 | RHO GTPase Effectors | 6.35E-01 |
| 19 | R-HSA-5684996 | MAPK1/MAPK3 signaling | 6.87E-01 |
| 20 | R-HSA-1257604 | PIP3 activates AKT signaling | 6.87E-01 |
| 21 | R-HSA-6798695 | Neutrophil degranulation | 6.87E-01 |
| 22 | R-HSA-5683057 | MAPK family signaling cascades | 6.87E-01 |
| 23 | R-HSA-9006925 | Intracellular signaling by second messengers | 6.87E-01 |
| 24 | R-HSA-5688426 | Deubiquitination | 6.87E-01 |
| 25 | R-HSA-109582 | Hemostasis | 6.87E-01 |
| 26 | R-HSA-5673001 | RAF/MAP kinase cascade | 7.12E-01 |
| 27 | R-HSA-382551 | Transport of small molecules | 9.40E-01 |
| 28 | R-HSA-9006934 | Signaling by Receptor Tyrosine Kinases | 9.73E-01 |
| 29 | R-HSA-72163 | mRNA Splicing - Major Pathway | 9.73E-01 |
| 30 | R-HSA-69620 | Cell Cycle Checkpoints | 9.73E-01 |

**Supplementary Table 9.** Gene Ontology-based ORA result (Pediatric, IFN-γ vs IFN-γ + HMOs).

| Rank | ID | Description | Adjusted p-value |
|---|---|---|---|
| 1 | GO:0002181 | cytoplasmic translation | 2.05E-36 |
| 2 | GO:0042273 | ribosomal large subunit biogenesis | 1.25E-02 |
| 3 | GO:0042274 | ribosomal small subunit biogenesis | 1.25E-02 |
| 4 | GO:0042254 | ribosome biogenesis | 1.42E-02 |
| 5 | GO:0022613 | ribonucleoprotein complex biogenesis | 2.06E-02 |
| 6 | GO:0042255 | ribosome assembly | 2.06E-02 |
| 7 | GO:0000027 | ribosomal large subunit assembly | 8.24E-02 |
| 8 | GO:0006413 | translational initiation | 2.91E-01 |
| 9 | GO:0061615 | glycolytic process through fructose-6-phosphate | 2.97E-01 |
| 10 | GO:0006177 | GMP biosynthetic process | 3.02E-01 |
| 11 | GO:0032328 | alanine transport | 3.02E-01 |
| 12 | GO:0022618 | protein-RNA complex assembly | 4.53E-01 |
| 13 | GO:0031400 | negative regulation of protein modification process | 5.14E-01 |
| 14 | GO:0071826 | protein-RNA complex organization | 5.77E-01 |
| 15 | GO:1901652 | response to peptide | 5.77E-01 |
| 16 | GO:1901653 | cellular response to peptide | 5.77E-01 |
| 17 | GO:0009168 | purine ribonucleoside monophosphate biosynthetic process | 5.77E-01 |
| 18 | GO:0006417 | regulation of translation | 5.77E-01 |
| 19 | GO:0010952 | positive regulation of peptidase activity | 5.77E-01 |
| 20 | GO:0048872 | homeostasis of number of cells | 5.77E-01 |
| 21 | GO:0006919 | activation of cysteine-type endopeptidase activity involved in apoptotic process | 5.77E-01 |
| 22 | GO:0016072 | rRNA metabolic process | 5.77E-01 |
| 23 | GO:0006364 | rRNA processing | 5.77E-01 |
| 24 | GO:0015804 | neutral amino acid transport | 5.77E-01 |
| 25 | GO:0015824 | proline transport | 5.77E-01 |
| 26 | GO:0019852 | L-ascorbic acid metabolic process | 5.77E-01 |
| 27 | GO:0032261 | purine nucleotide salvage | 5.77E-01 |
| 28 | GO:1901334 | lactone metabolic process | 5.77E-01 |
| 29 | GO:0009127 | purine nucleoside monophosphate biosynthetic process | 5.77E-01 |
| 30 | GO:0051220 | cytoplasmic sequestering of protein | 5.77E-01 |

**Supplementary Table 10.** Gene Ontology-based GSEA result (Pediatric, IFN-γ vs IFN-γ + HMOs).

| Rank | ID | Description | Enrichment Score | NES | Adjusted p-value |
|---|---|---|---|---|---|
| 1 | GO:0002181 | cytoplasmic translation | -0.757408 | -3.328177 | 4.39E-38 |
| 2 | GO:0022613 | ribonucleoprotein complex biogenesis | -0.369795 | -1.850934 | 5.90E-07 |
| 3 | GO:0042254 | ribosome biogenesis | -0.391117 | -1.896365 | 1.48E-05 |
| 4 | GO:0042274 | ribosomal small subunit biogenesis | -0.534587 | -2.19041 | 3.05E-05 |
| 5 | GO:0042273 | ribosomal large subunit biogenesis | -0.569159 | -2.234999 | 1.23E-04 |
| 6 | GO:0071826 | protein-RNA complex organization | -0.422587 | -1.938839 | 1.23E-04 |
| 7 | GO:0022618 | protein-RNA complex assembly | -0.42454 | -1.935188 | 1.44E-04 |
| 8 | GO:0042255 | ribosome assembly | -0.544922 | -2.052412 | 7.91E-03 |
| 9 | GO:0006364 | rRNA processing | -0.364313 | -1.683607 | 7.98E-03 |
| 10 | GO:2001257 | regulation of cation channel activity | 0.619129 | 2.323298 | 2.41E-02 |
| 11 | GO:0000028 | ribosomal small subunit assembly | -0.749061 | -2.093904 | 5.85E-02 |
| 12 | GO:0000027 | ribosomal large subunit assembly | -0.656815 | -2.055892 | 7.41E-02 |
| 13 | GO:1901019 | regulation of calcium ion transmembrane transporter activity | 0.666775 | 2.3977 | 8.93E-02 |
| 14 | GO:0001732 | formation of cytoplasmic translation initiation complex | -0.725312 | -1.985795 | 9.38E-02 |
| 15 | GO:0048285 | organelle fission | 0.363096 | 1.714864 | 1.90E-01 |
| 16 | GO:0010638 | positive regulation of organelle organization | 0.351262 | 1.675518 | 1.90E-01 |
| 17 | GO:0044770 | cell cycle phase transition | 0.33218 | 1.587885 | 1.90E-01 |
| 18 | GO:0060314 | regulation of ryanodine-sensitive calcium-release channel activity | 0.828195 | 2.360429 | 2.20E-01 |
| 19 | GO:0032412 | regulation of monoatomic ion transmembrane transporter activity | 0.466777 | 1.918345 | 2.40E-01 |
| 20 | GO:0051279 | regulation of release of sequestered calcium ion into cytosol | 0.641785 | 2.325125 | 2.88E-01 |
| 21 | GO:0000413 | protein peptidyl-prolyl isomerization | 0.789714 | 2.250755 | 2.88E-01 |
| 22 | GO:0031112 | positive regulation of microtubule polymerization or depolymerization | 0.653879 | 2.098555 | 2.88E-01 |
| 23 | GO:0090311 | regulation of protein deacetylation | 0.649846 | 2.085612 | 2.88E-01 |
| 24 | GO:0050848 | regulation of calcium-mediated signaling | 0.577679 | 2.077314 | 2.88E-01 |
| 25 | GO:1903169 | regulation of calcium ion transmembrane transport | 0.517182 | 2.069978 | 2.88E-01 |
| 26 | GO:0010288 | response to lead ion | 0.754829 | 2.062148 | 2.88E-01 |
| 27 | GO:0010917 | negative regulation of mitochondrial membrane potential | 0.835133 | 2.060373 | 2.88E-01 |
| 28 | GO:0045837 | negative regulation of membrane potential | 0.835133 | 2.060373 | 2.88E-01 |
| 29 | GO:0031116 | positive regulation of microtubule polymerization | 0.652218 | 2.059996 | 2.88E-01 |
| 30 | GO:0048311 | mitochondrion distribution | 0.794868 | 2.055634 | 2.88E-01 |

**Supplementary Table 11.** Inflammatory genes whose expression changes caused by IFN-γ are mitigated by pre-treatment with HMOs.

| | | Pediatric, NT vs IFN-γ | | Pediatric, IFN-γ vs IFN-γ + HMOs | |
|---|---|---|---|---|---|
| **Gene id** | **Symbol** | **$Log_2$Fold-Change** | **Adjusted p-value** | **$Log_2$Fold-Change** | **Adjusted p-value** |
| ENSG00000142192 | APP | -0.321321 | 1.87E-09 | 0.147944 | 0.039247 |
| ENSG00000171150 | SOCS5 | -0.308844 | 2.86E-04 | 0.260317 | 0.003443 |
| ENSG00000176903 | PNMA1 | -0.199017 | 4.70E-02 | 0.28343 | 0.003508 |
| ENSG00000050426 | LETMD1 | 0.296549 | 3.78E-04 | -0.19783 | 0.032904 |
| ENSG00000148344 | PTGES | 0.413192 | 3.14E-02 | -0.198356 | 0.032342 |
| ENSG00000118503 | TNFAIP3 | 0.355691 | 5.08E-03 | 0.208887 | 0.041139 |
